# A common neuronal basis for Pavlovian and instrumental learning in amygdala circuits

**DOI:** 10.1101/2025.01.13.632684

**Authors:** Coline Riffault, Yael Bitterman, Sigrid Müller, Andreas Lüthi, Julien Courtin

**Affiliations:** Neurocentre Magendie, Inserm U1215, Bordeaux, France; University of Bordeaux, Bordeaux, France; Friedrich Miescher Institute for Biomedical Research, Basel, Switzerland; The Hebrew University of Jerusalem, Ein Kerem, Jerusalem, Israel

**Author notes:** These authors contributed equally (Andreas Lüthi, Julien Courtin).

## Abstract

Reward-predictive Pavlovian cues can selectively invigorate instrumental behaviors. Accumulating evidence pinpoints the basolateral amygdala (BLA) as a key brain structure to assign outcome-specific motivational significance to Pavlovian cues or Instrumental actions. However, whether neuronal representations of Pavlovian and Instrumental learnings are processed in different BLA circuits and how the resulting memories interact is unknown. Here, we used calcium imaging and optogenetic manipulation of the BLA neurons in mice that acquired and expressed Pavlovian and Instrumental behaviors in a multi-phase behavioral task called specific Pavlovian to Instrumental Transfer (sPIT). We first confirmed that Pavlovian cues selectively invigorate instrumental actions and showed that this effect, referred as the sPIT effect, depends on BLA activity. Then, by tracking the activity of single BLA neurons across days, we found that BLA neurons integrate common reward-anticipation information relevant for Pavlovian and instrumental learning. Interestingly, this reward-anticipation information re-emerged during subsequent memory transfer and covaried with sPIT effect. These findings reveal that the BLA encodes an outcome-specific motivational state that generalizes across Pavlovian and instrumental behaviors to promote and guide reward-seeking behavior.

## INTRODUCTION

Environmental cues have the property to influence ongoing behavior^1^. This phenomenon can be readily examined by using the Pavlovian to Instrumental Transfer (PIT) paradigm^2^, in which the subject is sequentially exposed to Pavlovian and instrumental conditioning followed by a PIT test. PIT studies have shown that rewarding or aversive features of Pavlovian stimuli can affect the expression of ongoing instrumental behavior. Behavioral^3, 4^ and computational^5, 6^ research has led to the general concept that, in the context of appetitive PIT, a reward-predictive Pavlovian stimulus produces an outcome expectancy which then biases the instrumental behavior leading to that reward^7^. This phenomenon implies that dedicated brain circuits are able to encode specific features of this unique outcome, during both Pavlovian and instrumental learnings.

In support of this, BLA has been shown to encode both appetitive Pavlovian and instrumental learning^8, 9^. In particular, it has been shown that BLA neurons assign specific motivational significance to Pavlovian stimuli^10^ or instrumental actions^11^ that predict appetitive outcomes. Moreover, lesion and inactivation experiments pinpoint the BLA as a key brain structure supporting the outcome-specific form of PIT, i.e., sPIT^12-14^. However, whether encoding of Pavlovian and instrumental information is supported by shared or separate BLA circuits is unknown.

## RESULTS

### Pavlovian cues selectively invigorate instrumental actions

To address the neuronal basis of sPIT in the BLA, we used 1 photon calcium imaging in freely moving mice to record the neuronal activity of the main population of BLA neurons, i.e., the Calcium/calmodulin-dependent protein kinase 2 (CaMK2)-expressing principal neurons across the different phases of a sPIT paradigm specifically adapted to the application of optogenetic and calcium imaging approaches. Food-restricted male mice were sequentially exposed to Pavlovian conditioning (PavC), instrumental conditioning (InsC) and a final sPIT test (Fig. 1A). Pavlovian and instrumental contingencies were learned independently in different contexts but involved the same rewards (20% sucrose or sweet milk liquid solutions). During the four days of Pavlovian training, two different tones (conditioned stimuli: CS1 and CS2) were paired with two different rewards (30 µl of 20% sucrose or sweet milk liquid solutions, unconditioned stimuli: US1 and US2). By day 4 of training, mice showed robust anticipatory licking, i.e., a Pavlovian conditioned response during the CSs (Fig. 1B). Mice were then exposed to five days of instrumental training, during which one of two actions (action1 and action2, lever press) led to the delivery of the sucrose reward and the other to the delivery of the milk reward (20 µl of reward 1 and reward2), with no explicit stimuli signalling trial start or reward availability. As previously described^11^, by day 5 of training, mice displayed high goal-directed action performance (Fig. 1B). Finally, the mice were exposed to the sPIT session during which both actions were available. Importantly, no rewards were delivered during the sPIT session. As a first phase, instrumental action responses were extinguished for 12 min (pre-CS period) to reduce the baseline action rate (Fig. 1C-D). Then, 4 CS1 and 4 CS2 were presented to the mice always in the same order (Fig. 1C; CSs were the same than Pavlovian CSs). During this second phase (CS period), mice exhibited clear evidence of sPIT effect. Indeed, mice showed a significant and selective increase of instrumental action performance during CS presentations. Specifically, a CS motivates and biases the performance and choice of the specific action that earns the same outcome as that it predicts (e.g., CS1 invigorates action1 performance; Fig. 1C, E; Fig. S1A-B). Remarkably, CS presentations significantly increased the time spent by mice close to the lever zones without increasing licking behavior. This suggests that CS exposure preferentially promote instrumental actions rather than a general reward-seeking behavior (Fig. S1). To further validate our sPIT behavioral paradigm, characterized by a short Pavlovian training and relatively short CSs in comparison to classical paradigms^15-17^, we assessed whether the observed sPIT effect was sensitive to changes in the associative properties of Pavlovian CSs by performing a CS-US unpaired experiment. This manipulation of CS-US contingency abolished sPIT effect demonstrating that, in our conditions, CSs acquire sufficient motivational significance to selectively invigorate instrumental actions without affecting other behaviors (Fig. 1B-E; Fig. S1C-D).

**Figure 1:**
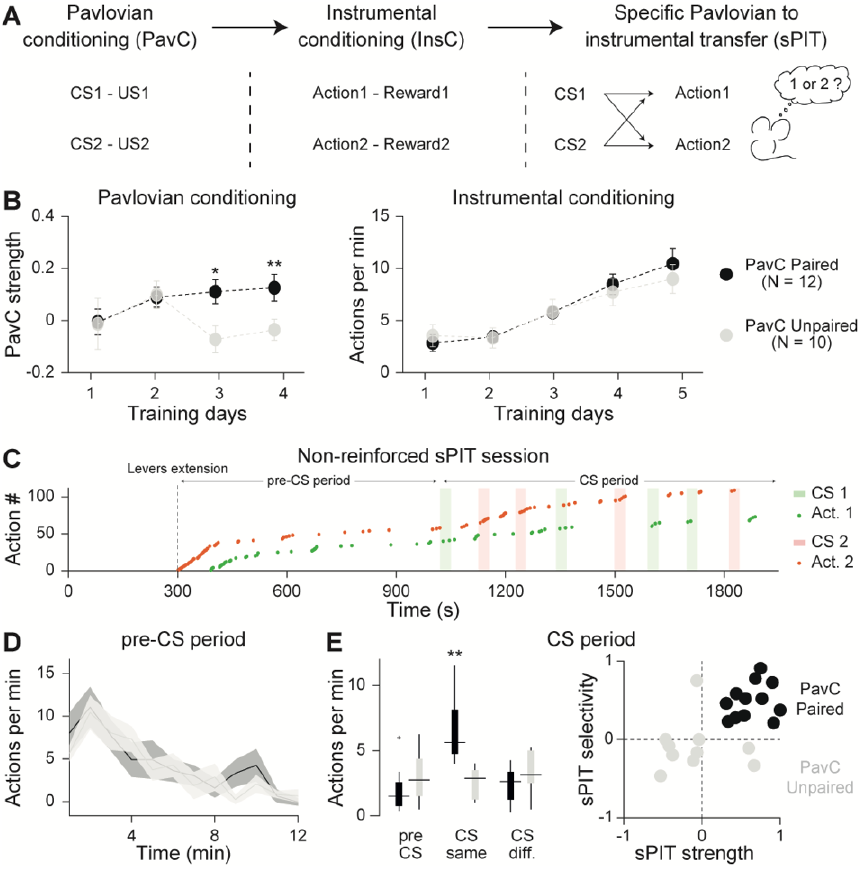
Pavlovian cues selectively invigorate instrumental actions. (**A**) Experimental design. During Pavlovian conditioning (PavC), 12 CS1 and 12 CS2 were presented (30 seconds tone stimuli; 4 kHz and 10 kHz) respectively paired with US1 and US2: 30 µl of sucrose or milk reward. During instrumental conditioning (InsC), action1 led to the delivery of 20 µl of sucrose reward and action2 to the delivery of 20 µl of milk reward. Action availability is controlled and the required number of actions per outcome increases over training days from continuous reinforcement to variable ratio 5. During specific Pavlovian to instrumental transfer (sPIT), 4 CS1 and 4 CS2 were presented while mice were free to perform action1 or action2. (**B**) PavC strength (see Methods) over Pavlovian training days (left) and number of actions per minute over instrumental training days (right) for PavC paired group (black, N = 12 mice) and unpaired group (gray, N = 10 mice; two-sided Wilcoxon rank-sum test comparing inter-groups PavC strength, *P < 0.05, **P < 0.01). Circles represent mean and error bars denote s.e.m.. (**C**) Structure of a non-reinforced sPIT session for one mouse. Vertical dashed line: lever extension and initiation of the 12 minutes of pre-CS period. During CS period, green and red areas: CS1 and CS2 (conserved order across mice); green and red dots: individual actions. (**D**) Action per minute during pre-CS period. Lines represent mean and shadings denote s.e.m.. (**E**) Left, action per minute before CS (pre CS) and during CSs for which CS and action outcome-identity match (CS same: PavC CS and InsC action associated with the same reward: action1 during CS1 or action2 during CS2) or differ (CS diff.: action1 during CS2 or action2 during CS1) for PavC paired and unpaired groups (two-sided Wilcoxon signed-rank and rank-sum tests comparing intra- and inter-groups, **P < 0.01). Box-and-whisker plots indicate median, interquartile, extreme data values and outliers of the data distribution. Right, sPIT strength and sPIT selectivity ratios (see Methods) showing individual sPIT effect. The sPIT effect was clear for the PavC paired group and abolished for the PavC unpaired group (two-sided Wilcoxon signed-rank and rank-sum tests comparing intra- and inter-groups, P < 0.01 for the two ratios).

### sPIT effect requires BLA activity

To test whether the sPIT effect depends on BLA activity, we optogenetically inhibited the activity of BLA CaMK2-expressing principal neurons (PNs) during CS presentations of the sPIT session, using an optogenetic approach. We virally expressed archaerhodopsin (ArchT) or GFP (control group) in BLA PNs and delivered light (589 nm) from CS onset until CS offset for half of the CS presentations in a pseudo-randomized manner. We first confirmed the efficiency of this approach to inhibit BLA PNs by combining optogenetic inhibition with electrophysiological single unit recordings. Putative BLA PNs were inhibited during the period of light stimulation (30 seconds; Fig. 2B). Mice expressing ArchT or GFP were then subjected to Pavlovian and instrumental conditioning (Fig. 2A; Fig. S2B). During the sPIT session, we quantified and compared the sPIT effect for CSs with and without light delivery over the BLA (Laser-ON versus Laser-OFF). Optogenetic inhibition of BLA PNs during CS presentations completely abolished the sPIT effect (Fig. 2C; Fig. S2C), demonstrating the necessity of acute BLA PN activity for sPIT effect.

**Figure 2:**
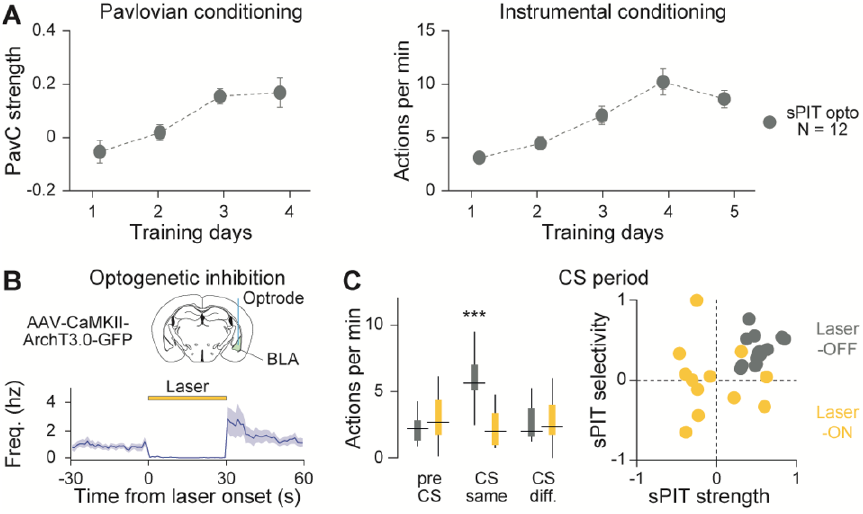
BLA PNs activity is necessary for sPIT effect. (**A**) PavC strength (see Methods) over Pavlovian training days (left) and number of actions per minute over instrumental training days (right) for optogenetic group (N = 12 mice). Circles represent mean and error bars denote s.e.m.. (**B**) Optogenetic inhibition of BLA PNs using archaerhodopsin (ArchT3.0) genetically expressed in CaMKII neurons (top). PSTH showing mean activity changes for putative PNs upon yellow light (bottom; laser-ON for 30 seconds; 20 repetitions; N = 2 mice; n = 14 putative PNs). Lines represent mean and shadings denote s.e.m.. (**C**) Left, action per minute before CS (pre CS) and during CSs for which CS and action identity match (CS same: PavC CS and InsC action associated with the same reward: action1 during CS1 or action2 during CS2) or differ (CS diff.: action1 during CS2 or action2 during CS1) for CSs without light (dark gray, laser-OFF) and with light (yellow, laser-ON) for ArchT mice (two-sided Wilcoxon signed-rank and rank-sum tests comparing intra- and inter-groups, ***P < 0.001). Box-and-whisker plots indicate median, interquartile, extreme data values and outliers of the data distribution. Right, sPIT strength and sPIT selectivity ratios (see Methods) for laser-OFF and laser-ON CSs in ArchT mice. The sPIT effect was abolished by optogenetic inhibition of BLA PNs activity (two-sided Wilcoxon signed-rank and rank-sum tests comparing intra- and inter-groups, P < 0.05 for the two ratios).

### Shared BLA neuronal ensembles for learning and sPIT

To understand which aspects of sPIT brain mechanisms are encoded by BLA PNs, we tracked BLA single neuron and neuronal ensemble activity across the entire sPIT paradigm using deep brain calcium imaging of BLA PNs with a miniaturized microscope and GCaMP6f as a calcium indicator^11^ (Fig. S3A). Like untethered mice (Fig. 1), the mice carrying le miniscope learned Pavlovian and instrumental conditioning and displayed a strong sPIT effect (Fig. S3B-C). We then extracted and analyzed the activity of single BLA PNs that could be registered from the last day of PavC (day 4), to the last day of InsC (day 5) and to the sPIT session (see Methods; 365 neurons from 9 mice; Fig. S3D). We first independently characterized functional types of neurons, for each session and for each CS, US, action and outcome showing a significant and consistent increase or decrease in activity at behaviorally relevant time points^11, 18-21^: CS onsets for PavC and sPIT, USs consumption onset for PavC, and both action and outcome consumption period onsets for InsC. During PavC, we identified four functional types of neurons based on distinct responses to CSs and USs. PavC CS^up^ neurons (n = 37/365) and PavC US^up^ neurons (n = 55/365) displayed increased activity during PavC CS presentations and PavC US consumption, respectively (Fig. 3A). CS^up^ and US^up^ neurons were not overlapping, suggesting a segregated CS and US information encoding (χ^2^ test, P <0.001). An equivalent proportion of neurons showed decreased activity during PavC CS presentations, the PavC CS^down^ neurons (n = 41/365) and during PavC US consumption, the PavC US^down^ neurons (n = 54/365; Fig. S4A). As for stimuli-activated neurons, CS^down^ and US^down^ neurons were not overlapping (χ^2^ test, P <0.001). Overall, neurons showing activity changes during behaviourally significant PavC times represented about 41% of the recorded neurons (n = 151/365). Remarkably, in contrast to the other ensembles, a significant proportion of PavC CS^up^ neurons showed no CS-selectivity, i.e., responding similarly to both to CS1 and CS2 (Fig. 3A, Fig. S4A; χ^2^ test, P = 0.96). We controlled for this observation by assessing the CS-selectivity with a larger number of CS^up^ neurons by using the single session dataset (for PavC session; n = 35 CS1-, 45 CS2-, 37 CS1CS2-responsive neurons; χ^2^ test, P = 0.35). During InsC, as previously described^11^, we identified two neuronal ensembles. InsC action neurons (n = 47/365) were activated at action period onsets and InsC consumption neurons (n = 60/365) exhibited increased activity time locked to rewarded licks. These populations were characterized by opposite dynamics and strong outcome-selectivity (Fig. 3B; action: χ^2^ test, P < 0.001; consumption: χ^2^ test, P < 0.05). Overall, InsC functional cell types represented about 29% of the recorded neurons (n = 107/365). During sPIT test, we identified two neuronal ensembles. sPIT CS^up^ neurons (n = 59/365) and sPIT CS^down^ neurons (n = 65/365) exhibited increased and decreased activity during sPIT CS presentations, respectively (Fig. 3C, Fig. S4B). Only the CS^up^ neurons were CS-selective (χ^2^ test, P = 0.058). Overall, sPIT functional ensembles represented about 34% (n = 124/365) of the recorded neurons.

**Figure 3:**
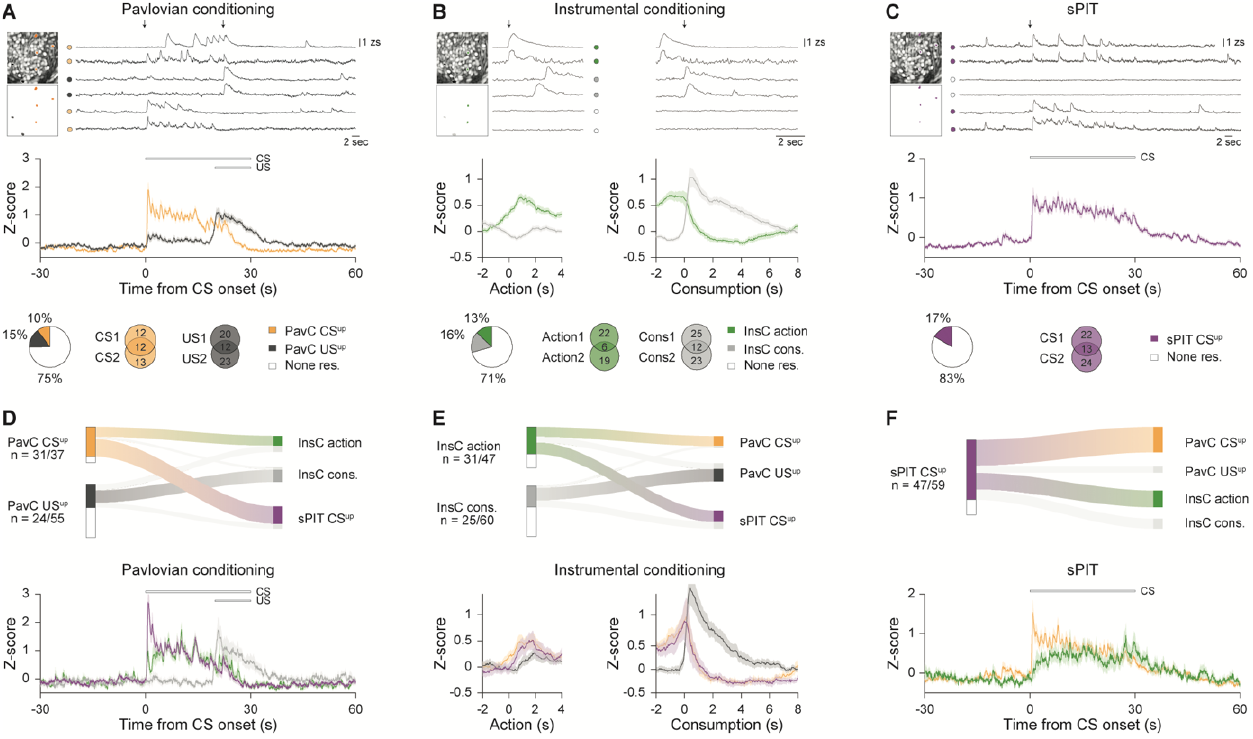
Pavlovian and instrumental neuronal ensembles re-emerge during sPIT. (**A**) Top, examples of spatial location within the field of view and calcium traces of six simultaneously recorded individual neurons during the last day of Pavlovian conditioning (colors indicate functional groups and black arrows CS and US onsets). Middle, average responses for PavC functional groups (orange: n = 37 PavC CS^up^ neurons; dark gray: n = 55 PavC US^up^ neurons) aligned to CS onsets (N = 9 mice, n = 365 neurons tracked across targeted behavioral sessions). Lines represent mean and shadings denote s.e.m.. Bottom, proportion (left) and CSs and USs selectivity (right; χ^2^ test, CS^up^: P = 0.96; χ^2^ test, US^up^: P = 0.058) of the PavC-modulated neurons. (**B**) As in panel A, examples, average responses, proportion and action and reward consumption selectivity (action: χ^2^ test, P < 0.001; consumption: χ^2^ test, P < 0.05) for InsC functional groups recorded during the last day of instrumental conditioning (green: n = 47 InsC action neurons; light gray: n = 60 InsC consumption neurons; N = 9 mice). Lines represent mean and shadings denote s.e.m.. (**C**) As in panels A and B, examples, average responses, proportion and CSs selectivity (χ^2^ test, P = 0.058) for sPIT functional group recorded during sPIT test (purple: n = 59 sPIT CS^up^ neurons; N = 9 mice). Lines represent mean and shadings denote s.e.m.. (**D**) Top, sankey plot illustrating the relationships of PavC functional groups with InsC and sPIT functional groups (n = 31/37 of PavC CS^up^ belong to InsC and sPIT groups; n = 24/55 of PavC US^up^). Colored paths denote significant overlaps between functional groups (P < 0.01, permutation test). Bottom, average responses aligned to CS onsets for overlapping neurons (green: n = 10 InsC action neurons; ligth gray: n = 13 InsC consumption neurons; purple: n = 18 sPIT CS^up^ neurons). (**E**) Top, sankey plot illustrating the relationships of InsC functional groups with PavC and sPIT functional groups (colored paths: P < 0.01, permutation test). Bottom, average responses aligned to action and consumption periods onsets for overlapping neurons (orange: n = 10 PavC CS^up^ neurons; dark gray: n = 13 PavC US^up^ neurons; purple: n = 15 sPIT CS^up^ neurons). (**F**) Top, sankey plot illustrating the relationships of sPIT functional groups with PavC and InsC functional groups (colored paths: P < 0.01, permutation test). Bottom, average responses aligned to CS onsets for overlapping neurons (orange: n = 18 PavC CS^up^ neurons; green: n = 15 InsC action neurons).

Next, we examined the degree of overlap across session between the different neuronal ensembles. In total, about 38% of the neurons (n = 137/365) belonged to neuronal ensembles of different sessions suggesting that, distinct task-features were integrated at single neuron level. We then determined for each pair of neuronal ensembles their overlaps across session (Fig. S3E). First, by comparing neuronal population activity across the two conditioning sessions, we identified a significant overlap of PavC CS^up^ and InsC action neurons. Interestingly, these neurons displayed neuronal activity dynamics more similar to the one displayed by action neurons during InsC than CS^up^ neurons during PavC as the sharp CS onset response usually exhibited by CS^up^ was absent. Furthermore, we found an overlap between PavC US^up^ and InsC consumption neurons. These neurons showed similar neuronal activity dynamics characterized by a sustained increase of activity during reward consumption (Fig. 3D-E).

Considering all three sessions, we identified three additional overlaps between session-specific neuronal ensembles (Fig. 3D-F). Notably, both the PavC CS^up^ and InsC action neurons overlapped with the sPIT CS^up^ neurons. PavC CS^up^ and InsC action neurons were strongly re-activated during sPIT CS presentations and showed distinct activity dynamics. Indeed, PavC CS^up^ neurons exhibited a sharp CS onset response and InsC action neurons a slower response during CS presentations. By assessing the distribution of calcium transients, we confirmed that these activity dynamic features were not merely reflecting slow calcium dynamics (Fig. S3F). Finally, we found that PavC CS^down^ and sPIT CS^down^ neuronal populations overlapped and displayed similar activity dynamics. (Fig. S4C-D). Taken together, these results support the hypothesis that BLA PNs integrate common features of Pavlovian and instrumental conditioning relevant for to reward-seeking behavior and reward consumption. Furthermore, we showed that during sPIT CS presentations, both PavC CS and InsC action neurons re-emerged with activity dynamics reflecting the behavioural sequence characterizing the sPIT effect – CS invigorates action.

### Shared BLA activity patterns across learnings and sPIT

In addition to the single neuron analysis and to avoid putative biases inherent to the sub-selection of neurons, we investigated the activity dynamics at the neuronal population level. We obtained population activity vector by binning single neuron activity in 200 ms bins. We then tracked, for each session, the re-occurrence of population activity patterns during both conditioning episodes and associated with CS presentations, reward-seeking behavior and reward consumption. During PavC, CS onset related activity patterns (a in Fig. 4A) were maintained during the CS presentation until the start of US consumption. The same population activity pattern strongly re-occurred during sPIT CS presentation. PavC reward-anticipation pattern that occurred before US consumption (b in Fig. 4A), ramped from CS onset until the start of US consumption. This pattern re-occurred during action periods during InsC and during sPIT CS presentations similarly to the PavC CS pattern. Reward consumption patterns observed during US consumption (c in Fig. 4A), were restricted to the US consumption period during PavC and re-occurred during InsC action and consumption periods but not during sPIT CS presentations (Fig. 4A-C). During InsC, similar to the patterns observed during PavC, we differentiated between two reward-anticipation patterns (d and e in Fig. 4F), one occurring well before reward consumption onset during the action period (PavC CS pattern) and another pattern that occurred right before reward consumption onset (PavC reward-anticipation pattern). As previously described^11^, during InsC and upon re-exposure to the task (sPIT pre-CS period), these patterns were maintained for several seconds throughout the action period until reward consumption and extinguished under non-reinforced conditions (sPIT; Fig. 4F-G). During the other sessions, these patterns re-occurred during PavC and sPIT CS presentations without showing a clear difference in their temporal dynamics. Lastly, in contrast to PavC US consumption pattern, InsC reward consumption pattern was emerging only during reward consumption during InsC and PavC (Fig. 4E-G). As observed at the single neuron level, PavC CS and reward-anticipation representations strongly re-emerged during sPIT CS presentation. Importantly, reward-anticipation patterns were discriminating between sPIT CSs along their sPIT performance but not between CS identity. Indeed, these patterns were significantly more recurrent during sPIT CSs inducing a strong sPIT effect compared to CSs inducing a weak effect (Fig. 4D-H). Taken together, these results show that BLA neuronal activity patterns integrate information relevant for reward seeking behavior across Pavlovian and instrumental conditioning sessions and that this shared information mediates interactions between Pavlovian and instrumental memory systems.

**Figure 4:**
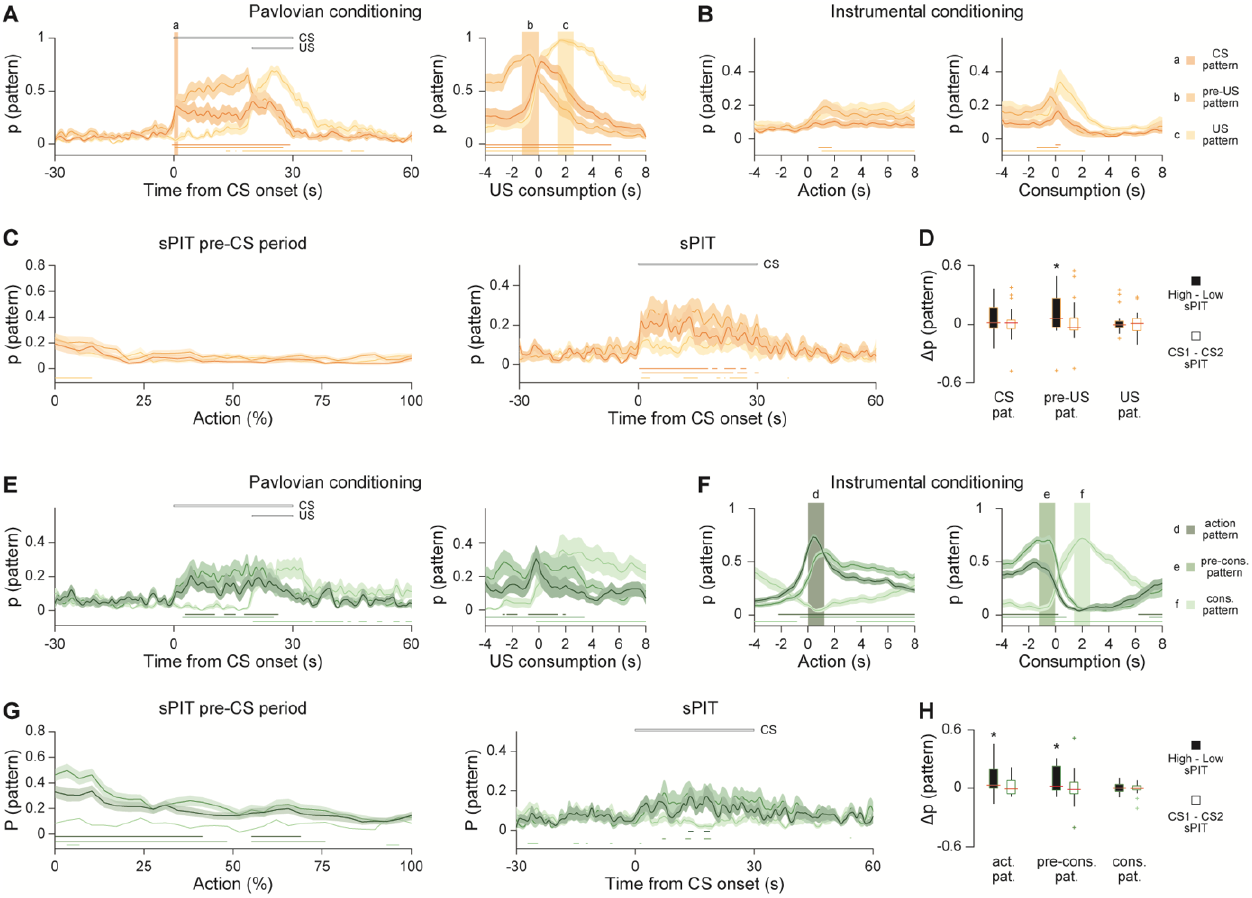
Shared neuronal population activity patterns across learnings and memory transfer. (**A**) Probability of PavC CS onset (darkest orange), PavC pre-US (intermediate orange) and PavC US (lighter orange) neuronal activity patterns to re-occur in response to CSs and USs during the last day of Pavlovian training. (**B**) Probability of PavC patterns to re-occur in response to action period onsets and reward consumption onsets during the last day of instrumental training. (**C**) Probability of PavC patterns to re-occur in response to action period onsets during pre-CS and CSs periods of sPIT test. Shaded areas denote reference population activity patterns. Lines represent mean and shadings denote s.e.m.. Significant pattern probabilities over average probability across the corresponding behavioral session are represented by horizontal lines (two-sided Wilcoxon signed-rank test, P < 0.05). (**D**) Difference in pattern probabilities for PavC patterns between CS high (CS with the highest sPIT effect) and CS low (CS with the lowest sPIT effect) to assess patterns action-dependency; and between CS1 and CS2 to assess patterns CS selectivity (two-sided Wilcoxon signed-rank test, *P < 0.05). (**E**) Probability of InsC action period onset (darkest green), InsC pre-consumption (pre-cons.; intermediate green) and InsC consumption (cons.; lighter green) neuronal activity patterns to re-occur in response to CSs and USs during the last day of Pavlovian training. (**F**) Probability of InsC patterns to re-occur in response to action period onsets and reward consumption onsets during the last day of instrumental training. (**G**) Probability of InsC patterns to re-occur in response to action period onsets during pre-CS and CSs periods of sPIT test. Shaded areas denote reference population activity patterns. Lines represent mean and shadings denote s.e.m.. Significant pattern probabilities over average probability across the corresponding behavioral session are represented by horizontal lines (two-sided Wilcoxon signed-rank test, P < 0.05). (**H**) Difference in pattern probabilities for InsC patterns between CS high and CS low; and between CS1 and CS2 (two-sided Wilcoxon signed-rank test, *P < 0.05).c

## DISCUSSION

Here we used deep-brain calcium imaging to monitor the activity dynamics of BLA PNs during key sessions of sPIT paradigm. We found that BLA encodes an outcome-specific motivational state that integrates outcome-specific Pavlovian and instrumental information during learning. This outcome-specific motivational state re-emerged during subsequent memory retrieval and transfer.

### BLA and Pavlovian and instrumental learning

Pavlovian and instrumental learning are often seen as two distinct phenomena, involving distinct neuronal mechanisms^4^. Quite in contrast to this notion, our results indicate that BLA single neuron and population activity reflect common variables, such as reward-seeking or reward consumption behavior, during Pavlovian and instrumental learning. Furthermore, BLA activity dynamics were stinkingly similar across Pavlovian and instrumental learning. In particular, the temporal relation of reward-anticipation patterns to reward consumption onset were conserved during Pavlovian and instrumental conditioning, suggesting common encoding principles.

### BLA and outcome-specific motivational state

Recent works highlight the role of the BLA in encoding behavioral states^11, 22, 23^. The present findings extend this concept. We show that, across the different behavioral sessions, neuronal ensembles and patterns can be sustained for several seconds during CS presentation or action period and that they eventually disappear at reward consumption onset. We recently characterized the outcome-specific motivational state occurring during instrumental actions^11^. Here, in another dataset, we confirm that this state emerges during instrumental action periods and that its maintenance necessitates reinforcement. We further showed that this state is emerging also during PavC and sPIT CS presentations supposably to motivate outcome-specific-seeking behavior.

### BLA and sPIT

Pre-sPIT test lesions or pharmacological inactivation of the BLA disrupt the expression of sPIT^12, 13^. Here we used temporally precise optogenetic approaches to show that inhibition of BLA PNs activity during sPIT CS presentations abolishes the sPIT effect. Furthermore, our recordings showed that BLA PNs displayed strong CS-evoked activity with distinct dynamics. Indeed, during sPIT CS presentations, neuronal ensembles and population activity patterns associated with PavC CS exposure preceded those associated with reward-seeking behavior, suggesting a causal relationship between them. Furthermore, patterns associated with reward-seeking behavior covaried positively with the sPIT effect. This result supports the hypothesis that during sPIT, the BLA is not simply relaying CS information but controlling instrumental actions. Overall, our data demonstrate that BLA activity patterns during Pavlovian and instrumental learning re-emerge during sPIT CS presentations. This strongly indicates that Pavlovian and instrumental memories share a common neuronal engram.

## METHODS

### Animals

All animal procedures were performed in accordance with institutional guidelines at the Friedrich Miescher Institute for Biomedical Research and were approved by the Veterinary Department of the Basel-Stadt Canton. Male mice (C57BL/6JRccHsd, Envigo) were used throughout the study. We chose to do experiments with male mice as they are on average heavier than female mice. This facilitates the carrying of the GRIN lenses implants during locomotion. Moreover, this allowed us to directly compare our present results to previous studies^11^. Mice were individually housed for at least 2 weeks before starting behavioral paradigms and were kept in a 12 h light–dark cycle. Mice well-being was monitored throughout the experimental period. All behavioral experiments were conducted during the light cycle.

### Surgical procedures

Surgical procedures were performed as previously described^11^. 8-week-old mice were anesthetized using isoflurane (3–5% for induction, 1–2% for maintenance; Attane, Provet) in oxygen-enriched air (Oxymat 3, Weinmann) and placed in a stereotactic frame (Model 1900, Kopf Instruments). Injections of buprenorphine (Temgesic, Indivior UK Limited; 0.1 mg per kg body weight subcutaneously 30 minutes before anesthesia) and ropivacain (Naropin, AstraZeneca; 0.1 ml locally under the scalp before incision) were provided for analgesia. Postoperative pain medication included buprenorphine (0.1 mg per kg in the drinking water; overnight) and injections of meloxicam (Metacam, Boehringer Ingelheim; 1 mg per kg subcutaneously) for 3 days after surgery. Ophthalmic gel was applied to avoid eye drying (Viscotears, Bausch+Lomb). The body temperature of the experimental animal was maintained at 36 °C using a feedback-controlled heating pad (FHC).

#### For deep-brain calcium imaging

AAV2/5.CaMK2.GCaMP6f (University of Zurich Vector Core) was unilaterally injected into the BLA (300 nl) using a precision micropositioner (Model 2650, Kopf Instruments) and pulled glass pipettes (tip diameter ∼20 μm) connected to a Picospritzer III microinjection system (Parker Hannifin Corporation) at the following coordinates from bregma: anterior–posterior (AP): -1.6 mm; medial–lateral (ML): -3.35 mm; dorsal–ventral (DV): 4.2 mm. After injection, a track above the imaging site was cut with a sterile needle to aid the insertion of the graded index lens (GRIN lens, 0.6 × 7.3 mm, Inscopix). The GRIN lens was subsequently lowered into the brain using a micropositioner through the track (4.3 mm from brain surface) with custom-built lens holder and fixed to the skull using ultraviolet light-curable glue (Henkel, Loctite 4305). Dental acrylic (Paladur, Heraeus) was used to seal the skull and attach a custom-made head bar for animal fixation during the miniature microscope mounting procedure. Mice were allowed to recover for 3 weeks after surgery before checking for GCaMP expression.

#### For optogenetic

AAV2/5.CaMK2.ArchT.GFP or AAV2/5.CaMK2.GFP (University of Zurich Vector Core) was bilaterally injected into the BLA (200 nl per hemisphere, coordinates from bregma: AP: - 1.6 mm; ML: -3.3 mm; DV: 4.2 mm). After injection, mice were bilaterally implanted with custom-made optic fiber connectors. Fiber tips were lowered to –3.9 mm below the brain surface using a micropositioner (Fig. S2A). Implants were fixed to the skull using cyanoacrylate glue (Ultra Gel, Henkel) and miniature screws (P.A. Precision Screws). Dental acrylic mixed with black paint (to avoid light spread) was used to seal the skull. Mice were allowed to recover for 3 weeks before behavioral training to ensure adequate virus expression.

### Behavioral context

#### Pavlovian conditioning behavioral context

The behavioral context used in this study measured 26 cm L × 25 cm W × 40 cm H and was enclosed inside an acoustic foam isolated box. The context was equipped with 2 custom-made lickports^11^ located at the middle of the wall, close to each other (4 cm apart). The 2 lickports were separated by a little barrier (to force mice to physically move in order to lick from one to another lickport). Each lickport allowed licking behavior monitoring and the delivery or collection of liquid rewards via remotely controlled syringe pumps (PHM-107, Med Associates) and vacuum. The context contained overhead speaker for delivery of auditory stimuli (Multi Field Speaker, Tucker Davis). Video camera recorded from above at 40 fps for video-tracking purposes using Cineplex Software (Cineplex, Plexon). All timestamps of camera frames, miniscope frames, lever presses and analog signals from lickports were recorded with Neural Recording Data Acquisition Processor system (at 40kHz, OmniPlex, Plexon). Behavior, optogenetic or miniscope were synchronized and controlled by a multi I/O processor (RZ6, Tucker Davis).

#### Instrumental conditioning and sPIT behavioral context

Instrumental conditioning and sPIT occurred in the same context. The context dimensions, lickports positions, speaker and data acquisition equipment were as in the Pavlovian conditioning context. The context was equipped with 2 custom-made lickports and 2 levers (left/right; ENV-312-2M, Med Associates). All *operanda* (2 instrumental actions apparatus, left/right; and 2 lickports, left/right) were located on the same wall. The 2 instrumental actions apparatus (left/right) were located at the extremes (5 cm from the wall limits). The levers edges were positioned 2 cm from the floor. Lever extensions/retractions were remotely controlled.

### Behavioral procedures

#### Food restriction

Mice were food-restricted to 85% of their free-feeding body weight 4 days before and throughout the behavioral experiments. Mice were fed about 2 hours after their daily behavioral sessions with about 2.5g of regular food.

#### Pavlovian conditioning

Pavlovian conditioning was preceded with a session in which either sucrose (20%) or sweetened condensed milk (15%, Régilait) solutions were accessible, each at a fixed lickport (right/left lickport respectively) upon licking (maximum duration 20 minutes or 20 of each outcome). Mice then started Pavlovian learning for 4 days. Pavlovian sessions were structured as follows: 1) 5 minutes without US available (licking behavior was possible but not rewarded). 2) 45 minutes during which 2 different pure tones (conditioned stimuli: CSs) were played and paired with the delivery of 2 different liquid rewards (unconditioned stimuli: USs). In total 12 CS1 and 12 CS2 (4 kHz and 10 kHz counterbalanced across mice, total duration of 30 s, consisting of 200 ms pips repeated at 0.8 Hz; 75 dB sound pressure level) were played with a pseudorandom inter-CS interval (range 60-120 seconds). CSs were played in blocks of 4 CSs (blocks order were counterbalanced). For PavC paired mice, 30 µl of USs were delivered 20 seconds after CS onsets and collected 5 seconds after CS offsets. For PavC unpaired mice, 30 µl of rewards were delivered for 15 seconds (as for paired condition) at pseudorandom times after CS offsets (range 20-60 seconds). 3) 2 minutes without CSs or USs.

#### Instrumental conditioning

Instrumental conditioning was performed as previously described^11^. Instrumental training started with constant reinforcement (CR) session, in which outcomes (20 µl of sucrose and milk) were delivered after each action performed by the mouse. In order to speed up learning, only during this day we disposed food onto levers. Mice then started instrumental learning with two days of CR (without food onto actions, termed day1 and day2). After CR sessions, mice went through variable action-outcome ratio training (VR), first with average ratio 3 on day 3 (VR3, between 1 and 5 following normal distribution, µ = 3, σ = 1.5) and 5 on day 4 and day 5 (VR5, between 1 and 11 following normal distribution, µ = 5, σ = 2.5). For CR and VR sessions outcome order was counterbalanced.

#### Specific Pavlovian to instrumental transfer

After Pavlovian and instrumental conditioning mice were exposed to a sPIT session (Fig. 1C), structured as follows: 1) 5 minutes without action or outcome availability. 2) 12 minutes during which the two actions (left and right levers) were available (pre-CS period). 3) 12 minutes during which the two actions were available and 4 CS1 and 4 CS2 were played (with same characteristics than Pavlovian CSs) in the following chronological order: CS1-CS2-CS2-CS1-CS2-CS1-CS1-CS2. 4) 2 minutes without action or outcome availability (OFF task). Importantly, sPIT session was non-reinforced.

### Behaviors analysis

We collected CS, US, action and lick timestamps together with mouse position tracking (Bonsai software^24^). To quantify Pavlovian conditioning, PavC strength ratio was computed for CS1 and CS2 (CS1 PavC strength = (nLicks^CS1^ – nLicks^preCS1^) / (nLicks^CS1^ + nLicks^preCS1^); nLicks^preCS^: number of licks during 20 seconds before CS; nLicks^CS^: number of licks during first 20 seconds of CS). For instrumental conditioning, action rate (actions per minute) was used as the quantifier of the instrumental performance. To quantify sPIT behavior, action rate (actions per minute) was calculated for three distinct periods for each action (action1 and action2). Pre-CS period: number of actions during 30 seconds before CS onsets; CS same and CS different (CS diff.): number of actions during 30 seconds after CS onsets. For CS same condition only the CSs and actions respectively predicting and leading to the same reward during Pavlovian and instrumental conditioning were considered (action1 during CS1 or action2 during CS2). For CS diff. condition CSs and actions of different identity were considered (action2 during CS1 or action1 during CS2). Individual sPIT effect was quantified for CS1 and CS2 by using sPIT strength and sPIT selectivity ratios (CS1 sPIT strength = (nAct^CS1^ – nAct^preCS1^) / (nAct^CS1^ + nAct^preCS1^); CS1 sPIT selectivity = (nAct^CS1^ – nAct^CS2^) / (nAct^CS1^ + nAct^CS2^); nAct^preCS^: number of actions during 30 seconds before CS; nAct^CS^: number of actions during first 30 seconds of CS). For calcium data analysis five type of timestamps were used: CS onsets (PavC and sPIT), USs consumption period onsets (PavC), action times (sPIT), action period onsets (InsC) and outcome consumption period onsets (InsC). Consumption period onsets were defined as the first lick bouts (series of two or more consecutive licks, with inter-lick interval less than 1 seconds) that occurred after an outcome delivery. Action period onsets was defined as the first action of the time from the first action until the last action between consumption periods (Fig. 1).

### Calcium imaging using the miniature microscope

#### Miniature microscope imaging

Four weeks after surgery, mice were head-fixed to check GCaMP expression using the miniature microscope (nVista HD, Inscopix Inc.). If the expression level was sufficient, mice were anaesthetized with isoflurane (3–5% for induction, 1–2% for maintenance; Attane, Provet) in oxygen-enriched air (Oxymat 3, Weinmann) to fix the miniature microscope baseplate (BLP-2, Inscopix) on top of the cranium implant, using blue-light curable glue (Vertise Flow, Kerr). The miniature microscope was then detached, the baseplate was capped with a baseplate cover (Inscopix) and the mouse was returned to its home cage. During the three days preceding instrumental training, mice were habituated to head-fixation (about 5 minutes) followed by a free exploration session in home cage (about 10 minutes). This daily procedure allowed mice to habituate to miniature microscope mounting and caring. It allowed us to select mice with high number of neurons and better signal quality (more than 50 neurons, N = 9 out of 16 mice included in the analysis) and to set the miniature microscope light-emitting diode (LED) power and electronic focus (for the nVista3 system). The miniature microscope was mounted immediately before each behavioral session by head-fixation. Imaging data was acquired using nVista HD software (Inscopix Inc.) at a frame rate of 20 Hz (exposure time 50 ms) with a LED power of 0.6–0.8 mW/mm^2^ (excitation irradiance at objective front surface) analog gain of 1 to 4 and a field of view of 1280×800 pixels window (about 768 × 480 µm, nVista3 system). Imaging frames timestamps were recorded with Neural Recording Data Acquisition Processor system (Plexon).

#### Motion correction

Imaging files produced by the nVista3 system were imported into Matlab 2017b, via custom code (https://github.com/fmi-basel/1Photon_Analysis). Two regions of interest (ROI) manually selected by the experimental users were used to correct for translational motion. To address background noise during motion correction, we subtracted a Gaussian blurred image on a frame-by-frame basis from the raw imaging data. We used FFT based Image registration^25^, to register the first 100 frames on one ROI, and then used the median image these 100 frames as a template to register the rest of the frames in the imaging session, until correction was lower than a user-defined threshold (typically less than 0.01 average pixel shift on average across all frames). The procedure was then repeated on the resulting motion corrected movie, using the second user-defined ROI, to ensure minimal non-rigid motion. We applied calculated shifts to the raw movie and used the raw, motion corrected movies for the extraction of calcium traces and all subsequent analysis.

#### Calcium traces extraction

We processed each session independently and used CNMF-E (Constrained Nonnegative Matrix Factorization for micro Endoscopic data), a recently developed algorithm based on non-negative matrix factorization^26^, for automatic extraction of calcium traces. We then excluded from the analysis ROIs based on anatomy (ROI size, shape, or vicinity to the edge of lens), low signal to noise ratio or large overlap in signal and spatial location with other neurons (> 60 % spatial overlap and >.6 Pearson’s correlation between the traces across the entire session). Linear trends across an entire session were removed from the calcium traces and further calculations were performed on the z-scores of the detrended traces.

#### Registration of neuron identities across imaging Session

After the individual sessions were extracted and sorted, taking day 5 as reference, we performed automatic neuron alignment between sessions using centroid and shape matching algorithms^27^. To ensure correct alignment visual inspection was performed to assess whether the imaged neuron was the same across days, the ROI shape and location in the FOV had to be consistent across sessions.

#### Histology

Following the completion of behavioral experiments, mice were deeply anaesthetized with urethane (2 g per kg of body weight; intraperitoneally) and transcardially perfused with PBS followed by 4% paraformaldehyde in PBS. Brains were removed and post-fixed overnight at 4°C. 80 μm coronal brain slices containing the BLA were cut with a vibratome (VT1000 S, Leica) and stored in PBS. Slices were washed for 10 minutes in PBS, given a 5 minutes exposure to 4′,6-diamidin-2-phenylindol (DAPI, 1:10,000, Sigma-Aldrich), and then washed 3x 15 minutes in PBS. Slices were mounted on glass slides, coverslipped and imaged using an Axioscan Z1 slide scanner (Carl Zeiss AG), equipped with a 10X air objective (Plan-Apochromat 10X/0.45). Mice were excluded post hoc if the GRIN lens was not placed in BLA (N = 2 mice excluded).

#### Imaging plan location

The position of the center of each GRIN lens was matched against a mouse brain atlas^29^.

### Optogenetic experiments

The experimenter was blinded to animal’s experimental cohort (virus condition, GFP or ArchT). Animals were randomly allocated to experimental groups and were later identified by unique markers for group assignment. Before behavioral experiments, all mice were habituated to the optical fiber connection procedure by handling and short head-restraining for at least 3 days. On the experimental days, implanted fibers (fiber numerical aperture of 0.48, fiber inner core diameter of 200 μm; Thorlabs) were connected to a custom-built laser bench (Life Imaging Services) with custom fiber patch cables. An acousto-optic modulator (AA Opto-Electronic) controlled the laser intensity (MGL-F-589, 589-nm wavelength, CNI Lasers). Laser power at the fiber tip was measured before every subject with an optical power and energy meter (PM100D, ThorLabs) and adjusted to reach an irradiance value of about 4 mW at fiber tip. To inhibit BLA PNs during CSs, the laser was switched on from CS onsets until CS offsets. Laser was controlled by a multi I/O processor (RZ6, Tucker Davis) and switched ON (laser-ON) for half of the CSs (Fig. 2). *Histology:* Mice were transcardially perfused (as described above) and optical fibers removed. Brains were post-fixed in 4% paraformaldehyde for at least 2 h at 4 °C and cut into 80-μm coronal slices using a vibratome (VT1000S). Sections containing the BLA were immediately mounted on glass slides and coverslipped. To verify the specificity of viral expression and fiber tip placement, sections were scanned with an Axioscan Z1 slide scanner (Carl Zeiss AG), equipped with a 10X air objective (Plan-Apochromat 10X/0.45). Fiber tip placements were matched against a mouse brain atlas^29^. Mice were excluded from the analysis post hoc if they did not show bilateral expression of the virus, if virus expression (cell bodies expressing GFP) was detected outside the BLA or if they did not exhibit correct fiber placement (<300 μm away from the BLA). A total of N 10 of 32 mice were excluded.

### Freely moving electrophysiology

#### Surgery

Two mice were unilaterally implanted with a custom-built optrode consisting of an optic fiber with an attached 16-wire electrode into the BLA using a micropositioner (coordinates from bregma: AP: -1.6 mm; ML: -3.3 mm; DV: 4.2 mm). Electrodes tips were cut at an angle to protrude approximately 300–500 μm beyond the fiber end. In this case, the surgical procedure was similar to the previous one described above in the optogenetic part. Implants were fixed to the skull using cyanoacrylate glue (Ultra Gel, Henkel) and miniature screws (P.A. Precision Screws). Dental acrylic was used to seal the skull. Mice were allowed to recover for 2 weeks before recordings started.

#### Single-unit recordings

Mice were habituated to headstages (Plexon) connection procedure by handling and short head-restraining for at least 3 days before experiments. The headstages were connected to a 32-channel preamplifier (gain ×100, band-pass filter 150 Hz to 9 kHz for unit activity and 0.7 Hz to 170 Hz for field potentials, Plexon). Spiking activity was digitized at 40 kHz and band-pass-filtered from 250 Hz to 8 kHz, and isolated by time-amplitude window discrimination and template matching using a Neural Recording Data Acquisition Processor system (OmniPlex, Plexon).

#### Single-unit analysis

Single-unit spike sorting was performed using Offline Spike Sorter software (Plexon). Principal component scores were calculated for unsorted waveforms and plotted in a three-dimensional principal component space; clusters containing similar valid waveforms were manually defined. A group of waveforms were considered to be generated from a single neuron if the waveforms formed a discrete, isolated, cluster in the principal component space and did not contain a refractory period of less than 1 ms, as assessed using autocorrelogram analyses. To avoid analysis of the same neuron recorded on different channels, we computed cross-correlation histograms. If a target neuron presented a peak of activity at a time that the reference neuron fired, only one of the two neurons was considered for further analysis. Putative inhibitory interneurons were separated from putative excitatory projection neurons using an unsupervised cluster algorithm based on Ward’s method. In brief, the Euclidian distance was calculated between all neuron pairs based on the three-dimensional space defined by each neuron’s average half-spike width (measured from trough to peak), the firing rate and the area under the hyperpolarization phase of the spike^28^. To assess the efficiency of our optogenetic setting we recorded the activity of BLA putative PNs while applying 30 seconds laser stimulations (20 repetitions).

### Data analysis

We used custom routines written in MATLAB (Mathworks) to perform all analyses.

#### Identification of tasks-modulated neurons

We identified independently for each CSs, USs, actions and outcomes, functional types of neurons showing significant and consistent increase or decrease activity at following behavioral times: CS onsets (PavC and sPIT), USs consumption period onsets (PavC), action period onsets (InsC) and outcome consumption period onsets (InsC). Z-scored traces were baselined to the entire session. For CS- and US-responsive PNs, we compared the activity before CS onsets (20 seconds for CS and 5 seconds for US) to the activity during CS or US (20 seconds for CS and 5 seconds for US) using the two-sided Wilcoxon signed-sum test. PNs with significant increased or decreased CS- or US-responses (P < 0.05) to at least 3 CSs or USs were considered as CS- or US-responsive. Following this strategy, we identified 6 distinct functional groups of PNs responding to CSs and USs. PavC CS^up^ and PavC CS^down^ as PNs respectively increasing and decreasing their activity during PavC CS presentations. PavC US^up^ and PavC US^down^ as PNs respectively increasing and decreasing their activity during PavC US presentations. sPIT CS^up^ and sPIT CS^down^ as PNs respectively increasing and decreasing their activity during sPIT CS presentations. For action- and consumption-responsive PNs we performed as previously described^11^. Briefly, neurons were classified as action- and consumption-responsive if the maximum of their average response in a reference time window fulfilled three criteria: 1) it exceeded a threshold on the maxima obtained for each neuron by a bootstrapping procedure using the same time window around random timestamps along the session (number of random timestamps matched the number of real timestamps, 1000 iterations, threshold set at the 99 percentile); 2) it exceeded the average response outside the restricted reference window; 3) it exceeded the upper limit of the 95% confidence interval around zero (a z-score value of 0), computed using student’s t-distribution with the empirical mean and standard error of the maxima of individual traces. This allowed us to capture InsC action and InsC consumption neurons (Fig. 3).

#### Overlap of neuronal ensembles across sessions

to determine whether the degree of overlap between neuronal ensembles across sessions we assessed whether the real number of neurons belonging to each ensembles exceeded a threshold number obtained by a bootstrapping procedure. For each pair of session and each pair of neuronal ensembles, we computed a threshold set at the 99 percentile (P < 0.01) of a distribution obtained by randomizing the neurons identities (ID from 1 to 365) from one session (10000 iterations with number of random neuron IDs matched the number of real neuron IDs).

#### Pairwise correlation between population activity patterns and probability of patterns to re-occur

We binned the single neuron activity in 200 msec bins to obtain a single population activity vector per time point, and quantified the similarity between two activity vectors by the pairwise Pearson’s correlation coefficient (r). To follow the re-occurrence of specific population activity patterns, we performed pairwise Pearson’s correlation between reference patterns (average of activity vectors within 1 second) and every single population activity vector within and across the three sessions. We then marked all bins with activity vectors that had a positive correlation with the reference to obtain the probability of pattern to re-occur (the lower bound of the 99% confidence interval of the Pearson’s correlation > 0). We tracked the dynamic of six different population activity patterns in response to CSs and USs during the last day of Pavlovian training, to actions and consumption periods during the last day of instrumental training and to actions and CSs during sPIT. The six patterns were: PavC CS (after CS onsets), PavC pre-US (before US consumption period onsets), PavC US (2 seconds after US consumption period onsets), InsC action period onset (after action period onsets), InsC pre-consumption (before InsC consumption period onsets) and InsC consumption (2 seconds after InsC consumption period onsets) (Fig. 4).

#### Statistics

Statistical analysis was performed with Matlab (Mathworks). Normality of the data was assessed using Shapiro-Wilk test and either parametric (paired or unpaired *t*-test) or non-parametric (Wilcoxon signed-rank or Wilcoxon rank-sum tests) tests were used. Box-and-whisker plots indicate median (vertical line), interquartile (horizontal thick line), extreme data values (horizontal thin line) and outliers (+ symbol) of the data distribution. No statistical methods were used to pre-determine sample sizes, but our sample sizes are similar to those generally employed in the field. Statistical tests are mentioned in the figure legends. ns indicate p-values higher than 0.05; *, ** and *** indicate p-values smaller than 0.05, 0.01 and 0.001, respectively.

## Data and code availability

Custom code and full datasets will be made available upon final publication and are currently available upon request from JC.

## Author contributions

JC designed the experiments. JC and CR performed the experiments with help from SM. JC and YB analysed behavior and deep-brain imaging data. JC and AL supervised the work. JC wrote the paper. All authors contributed to the interpretation of the data and commented on the manuscript.

## Acknowledgements

The authors thank all members of the Courtin and Lüthi labs for helpful discussions and comments. They thank Tobias Eichlisberger, Christian Müller and all staff of the FMI Animal Facilities for excellent technical assistance. They further thank the Facility for Imaging and Microscopy at the FMI.

## Supplementary Materials

Supplementary Figures 1-4

## Supplementary information

## SUPPLEMENTARY FIGURES

**Supplementary Figure 1:**
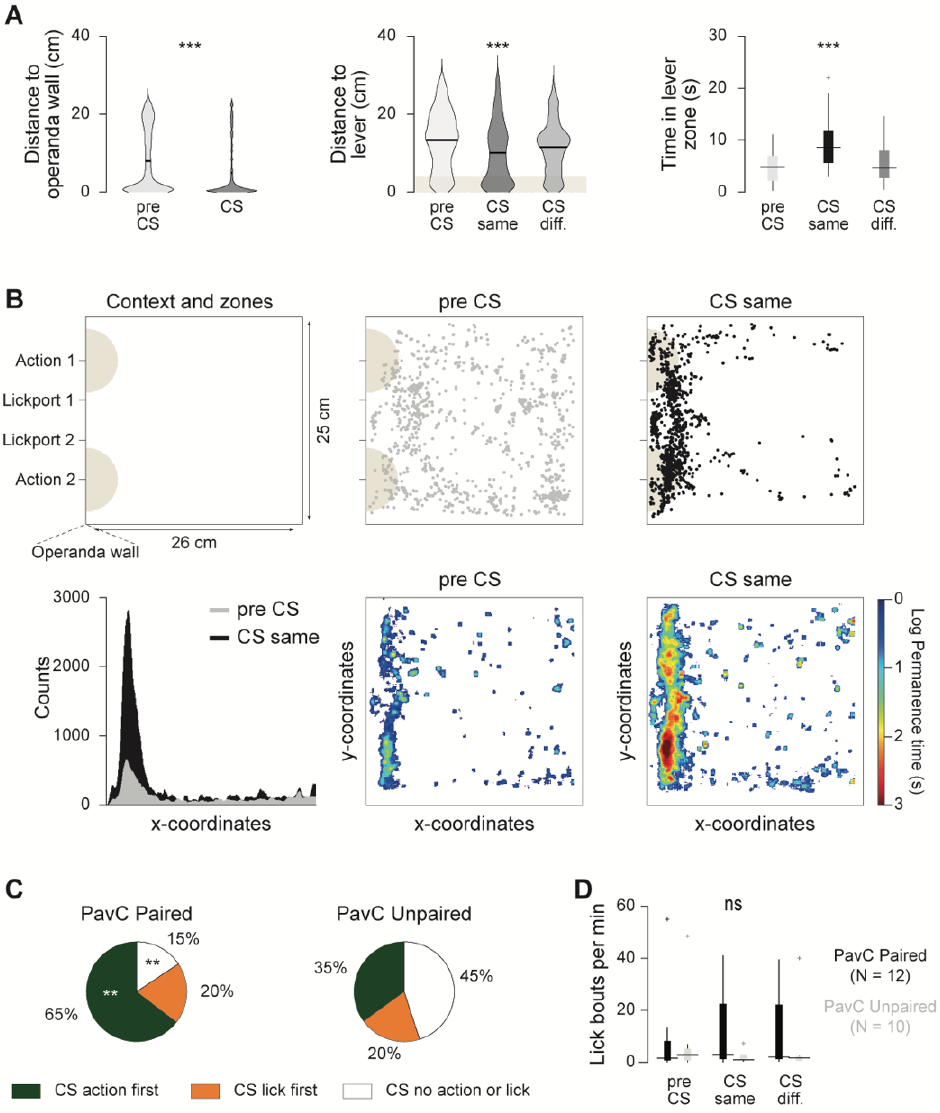
Pavlovian cues promote instrumental behaviors. (**A**) Distance between mice body center of mass and the operanda wall before and during CSs (left). Distance between mice body center of mass and the closest lever (middle) and time spend in lever zones (right; lever zone: 4 cm from lever) before CS (pre CS) and during CSs for which CS and action outcome-identity match (CS same: PavC CS and InsC action associated with the same reward: action1 during CS1 or action2 during CS2) or differ (CS diff.: action1 during CS2 or action2 during CS1) for PavC paired group (two-sided Wilcoxon signed-rank test, ***P < 0.001). Violin plots indicate median and the distribution of all data. Box-and-whisker plots indicate median, interquartile, extreme data values and outliers of the data distribution. (**B**) Context and zones dimensions and descriptions (top left). Trajectories of a mouse extracted from the video tracking before CS (top middle) and during CS same (top right). Mean time spend across the behavioral context before CS and during CS same for PavC paired mice (bottom). (**C**) Proportion of CSs during which mice performed first an instrumental action, a licking behavior or no behavior for PavC paired group (N = 12 mice) and unpaired group (N = 10 mice; two-sided Wilcoxon rank-sum test comparing inter-groups number of CS types, **P < 0.01). (**D**) Lick bouts per minute before CS (pre CS), during CS same and CS differ for PavC paired and unpaired groups (two-sided Wilcoxon signed-rank and rank-sum tests comparing intra- and inter-groups, P > 0.05). Box-and-whisker plots indicate median, interquartile, extreme data values and outliers of the data distribution.

**Supplementary Figure 2:**
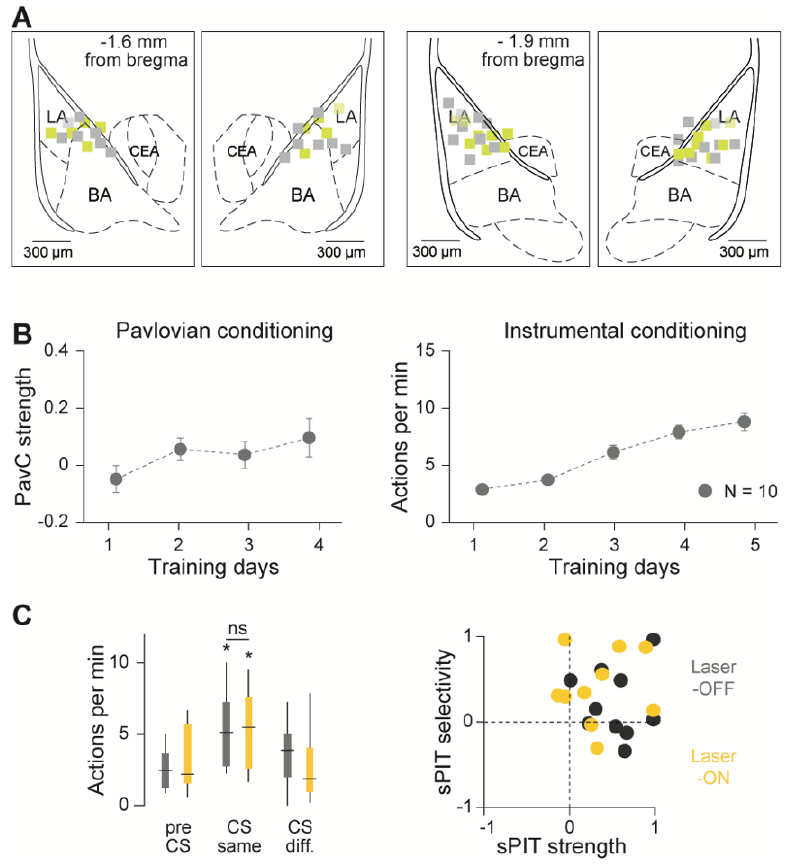
Controls of optogenetic inhibition experiments. (**A**) Anatomical position of optical fiber tips (gray: GFP mice, N = 10; yellow: ArchT mice, N = 12). (**B**) PavC strength (see Methods) over Pavlovian training days (left) and number of actions per minute over instrumental training days (right) for GFP group (N = 10 mice). Circles represent mean and error bars denote s.e.m.. (**C**) Left, action per minute before CS (pre CS) and during CSs for which CS and action identity match (CS same: PavC CS and InsC action associated with the same reward: action1 during CS1 or action2 during CS2) or differ (CS diff.: action1 during CS2 or action2 during CS1) for CSs without light (dark gray, laser-OFF) without light (yellow, laser-ON) for GFP control mice (two-sided Wilcoxon signed-rank and rank-sum tests comparing intra- and inter-groups, *P < 0.05). Box-and-whisker plots indicate median, interquartile, extreme data values and outliers of the data distribution. Right, sPIT strength and sPIT selectivity ratios (see Methods) for laser-OFF and laser-ON CSs in GFP mice. GFP control mice showed sPIT effect for CSs without and with laser.

**Supplementary Figure 3:**
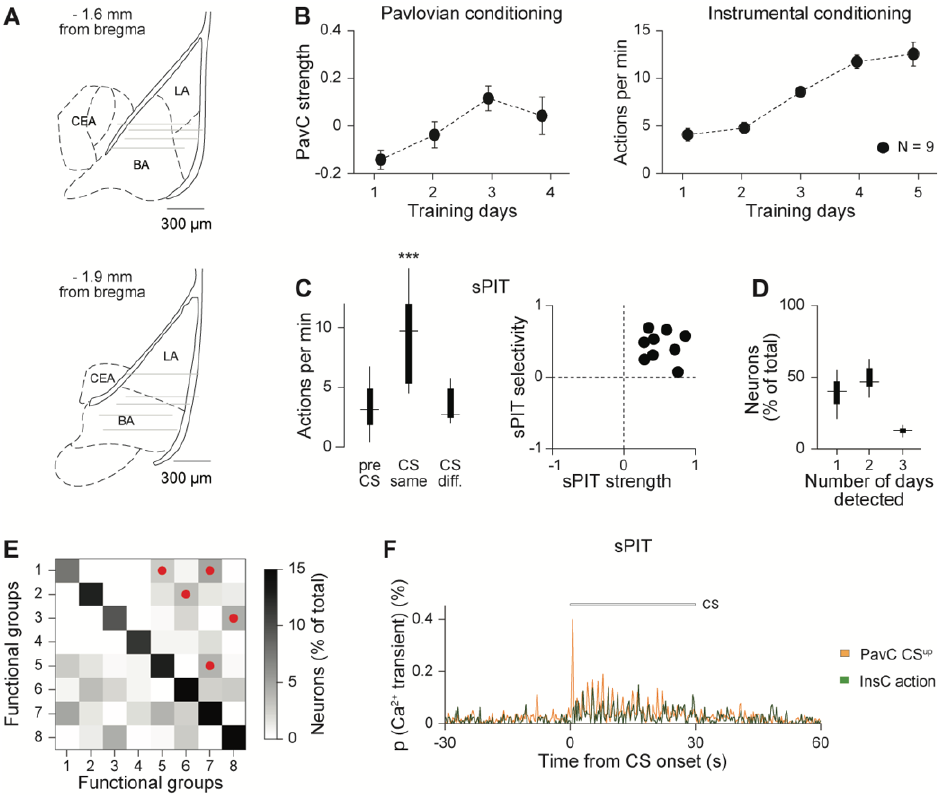
Tracking BLA PNs across days. (**A**) Anatomical location of the tip of the lenses for the 9 mice matched to a brain atlas. (**B**) PavC strength (see Methods) over Pavlovian training days (left) and number of actions per minute over instrumental training days (right) for miniscope mice (N = 9 mice). Circles represent mean and error bars denote s.e.m.. (**C**) Left, action per minute before CS (pre CS) and during CSs for which CS and action outcome-identity match (CS same: PavC CS and InsC action associated with the same reward: action1 during CS1 or action2 during CS2) or differ (CS diff.: action1 during CS2 or action2 during CS1) for miniscope mice (two-sided Wilcoxon signed-rank tests comparing intra-groups, ***P < 0.001). Box-and-whisker plots indicate median, interquartile, extreme data values and outliers of the data distribution. Right, sPIT strength and sPIT selectivity ratios (see Methods) showing individual sPIT effect. (**D**) Percentage of all 3266 neurons in the study that were detected in one of the three sessions. Box-and-whisker plots indicate median, interquartile, extreme data values and outliers of the data distribution. (**E**) Overlaps between PavC, InsC and sPIT functional groups (red dot: < 0.01, permutation test). (**F**) Calcium (Ca^2+^) transient probability of overlapping neurons in response to CSs during sPIT (as in Fig. 3F; orange: n = 18 PavC CS^up^ neurons; green: n = 15 InsC action neurons).

**Supplementary Figure 4:**
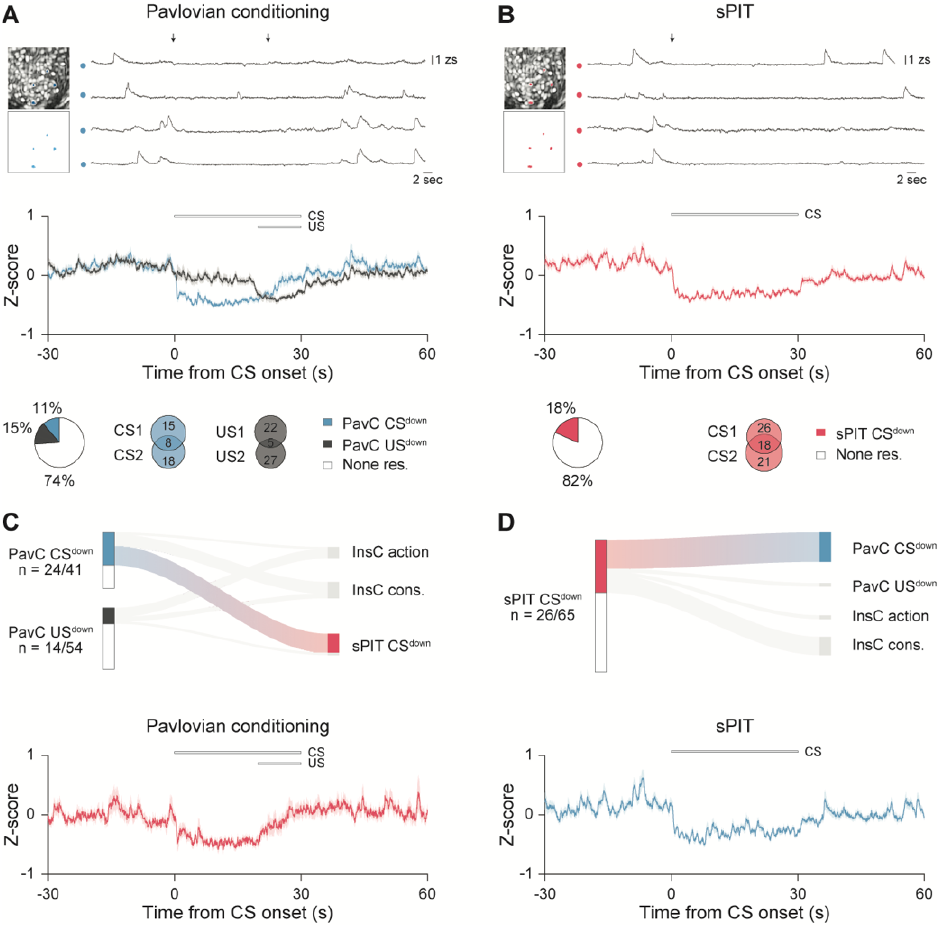
Pavlovian neuronal ensembles showing CS-evoked reduced activity re-emerge during sPIT CSs. (**A**) Top, examples of spatial location within the field of view (intensity correlation map) and calcium traces of four simultaneously recorded individual neurons during the last day of Pavlovian conditioning (colors indicate functional groups and black arrows CS and US onsets). Middle, average responses for PavC functional groups (blue: n = 41 PavC CS^down^ neurons; dark gray: n = 54 PavC US^down^ neurons) aligned to CS onsets (N = 9 mice, n = 365 neurons tracked across targeted behavioral sessions). Lines represent mean and shadings denote s.e.m.. Bottom, proportion (left) and CSs and USs selectivity (right: χ^2^ test, CS^down^: P < 0.05; χ^2^ test, US^down^: P <0.001) of the PavC-modulated neurons. (**B**) As in panels A, examples, average responses, proportion and CSs selectivity (χ^2^ test, P = 0.33) for sPIT functional group recorded during sPIT test (red: n = 65 sPIT CS^down^ neurons; N = 9 mice). Lines represent mean and shadings denote s.e.m.. (**C**) Top, sankey plot illustrating the relationships of PavC functional groups with InsC and sPIT functional groups. Colored paths denote significant overlaps between functional groups (P < 0.01, permutation test). Bottom, average responses aligned to CS onsets for overlapping neurons (red: n = 16 sPIT CS^down^ neurons). (**D**) Top, sankey plot illustrating the relationships of sPIT functional groups with PavC and InsC functional groups. Colored paths denote significant overlaps between functional groups (P < 0.01, permutation test). Bottom, average responses aligned to CS onsets for overlapping neurons (blue: n = 16 PavC CS^down^ neurons).

